# Deep tissue optical 3D imaging reveals preferential preservation of extra-islet β-cells in late-onset Type 1 Diabetes

**DOI:** 10.1101/2025.09.03.673632

**Authors:** Joakim Lehrstrand, Max Hahn, Björn Morén, Olle Korsgren, Tomas Alanentalo, Ulf Ahlgren

## Abstract

As residual β-cell function may have positive effects on diabetes regulation in T1D, details on its spatial distribution and mass could provide important information for potential future therapeutic regimens. We implemented an optical 3D imaging pipeline to generate a first account of the 3D-spatial and volumetric distribution of the remaining β-cells throughout the volume of an entire human late onset T1D pancreas at a microscopic resolution. As expected, β-cell mass was dramatically lower than in the non-diabetic pancreas. However, the pancreatic head displayed a morphology and size resembling the non-diabetic pancreas and had a 3 times higher β-cell density compared to the rest of the organ. Surprisingly, only a fraction of the residual β-cells were located within islet structures. Instead, the absolute majority were present as extra-islet β-cells, either as scattered individual cells or as clusters of β-cells, spatially separated from all other endocrine cell-types. Jointly, these extra-islet β-cells appeared roughly 60x less prone to succumb than islet associated β-cells. In sharp contrast to β-cell mass, α-cell density appeared unaffected. This 3D whole organ depiction of an entire long standing, late onset, T1D pancreas shows that individual β-cells may be preserved in a highly regionalized manner, potentially reflecting key aspects of disease dynamics.

## INTRODUCTION

Type 1 diabetes (T1D) is a chronic disease characterised by the progressive loss of pancreatic β-cells, ultimately leading to overt hyperglycaemia and a lifelong dependence on exogenous insulin therapy [1]. Despite extensive research, the exact aetiology remains unclear, however, evidence suggests that both genetic and environmental factors contribute to disease development, which results in a heterogenous patient group and the need for a personalised treatment approach[2].

An accumulating number of studies point to a link between age at T1D diagnosis and disease progression and severity. For example, histologically distinct T1D endotypes have been suggested depending on age of disease onset (under the age of 30) [3]. Further, people who are diagnosed later in life often retain a significant number of INS^+^ cells, despite being dependent on exogenous insulin [4] and refs. therein). Although T1D commonly has been considered a “juvenile” disease, several studies suggest that adult onset-T1D is more prevalent, and a high degree of late onset-T1D cases are misclassified as type 2 diabetes (T2D) due to overlapping clinical features[5].

Regardless of potential endotype or time of diagnosis, residual β-cell function may have positive effects on diabetes regulation and complications in T1D patients. This includes reduced risk of hyperglycaemia, improved metabolic control, lower exogenous insulin requirements, and lower risk of late complications. Therefore, understanding the spatial distribution, amount and function of residual β-cells could prove important for disease management. Whereas the presence of extra-islet β-cells, in addition to islet-associated β-cells, has been described in several publications ([6] and refs therein), the sheer size of the human pancreas (average range 71-83 cm^3^ [7]) and its complex organization, with numerous islets and/or β-cells scattered throughout the exocrine parenchyma [8], has effectively hindered detailed residual β-cell mass (BCM) assessments in T1D, especially in the whole organ context. Since access to larger tissue preparations of T1D pancreas, suitable for immunohistochemical analyses, is scarce, most studies aimed at surveying mass effects have been performed on limited tissue volumes, often by extrapolation of partial 2D data and/or with limited information about the tissués spatial origin within the gland.

Recently, we demonstrated an approach to generate a complete account of antibody labelled cells throughout the volume of the pancreas from deceased donors at microscopic resolution within a maintained spatial context. We could hereby lay bare the complete β-cell distribution in the non-diabetic (ND) pancreas, with information on the volume, shape and 3D coordinates for essentially every islet throughout the glandś volume [8]. In this report, we employed essentially the same imaging approach on an entire pancreas from a late onset-T1D (LO-T1D) donor. We demonstrate that the distribution of remaining β-cells could be highly heterogenous within the organ, and that islet associated β-cells may be around 60 times more susceptible to destruction than extra-islet β-cells. Hence, in sharp contrast to the ND pancreas, where extra-islet β-cells only constitute a fraction of the total BCM, extra-islet β-cells can several years after disease onset make up the absolute majority of the residual BCM in LO-T1D. Jointly, our results emphasize the value of a whole organ perspective in T1D studies and prompt for detailed investigations into the functional potential of extra islet-β-cells and their preservation.

## RESULTS

### 3D reconstruction of the residual β-cell mass distribution in LO-T1D displays regional differences in β-cell preservation

To assess the 3D-spatial distribution and volume of the residual β-cells mass in a LO-T1D donor (diagnosis at 50 years of age, disease duration 17 years, see **Table S1**, H2442), the pancreas was divided into tissue discs, using a 3D printed matrix (**Fig. S1**). These discs were antibody labelled and scanned individually by near infrared optical projection tomography (NIR-OPT) [9, 10]. Selected regions of interest (ROIs) were further examined by light sheet fluorescence microscopy (LSFM) [11, 12]. By aligning the resultant tomographic datasets together in 3D space, the entire pancreas could be reconstructed with regards to the 3D distribution of INS^+^ cells. Hereby, we generated a complete 3D data set of the remaining β-cell distribution throughout the volume of the gland (**Fig. 1** and **Movie S1**). Segmentation of INS^+^ signals provided a complete picture of all INS^+^ objects (for OPT data, an INS^+^ object is defined as a distinct body of INS^+^ cells that cannot be separated from each other at the resolution applied during segmentation, i.e., ∼21 µm, see methods) including their individual volumes and spatial 3D coordinates throughout the gland (For details see Methods and [8]).

**Figure 1.**
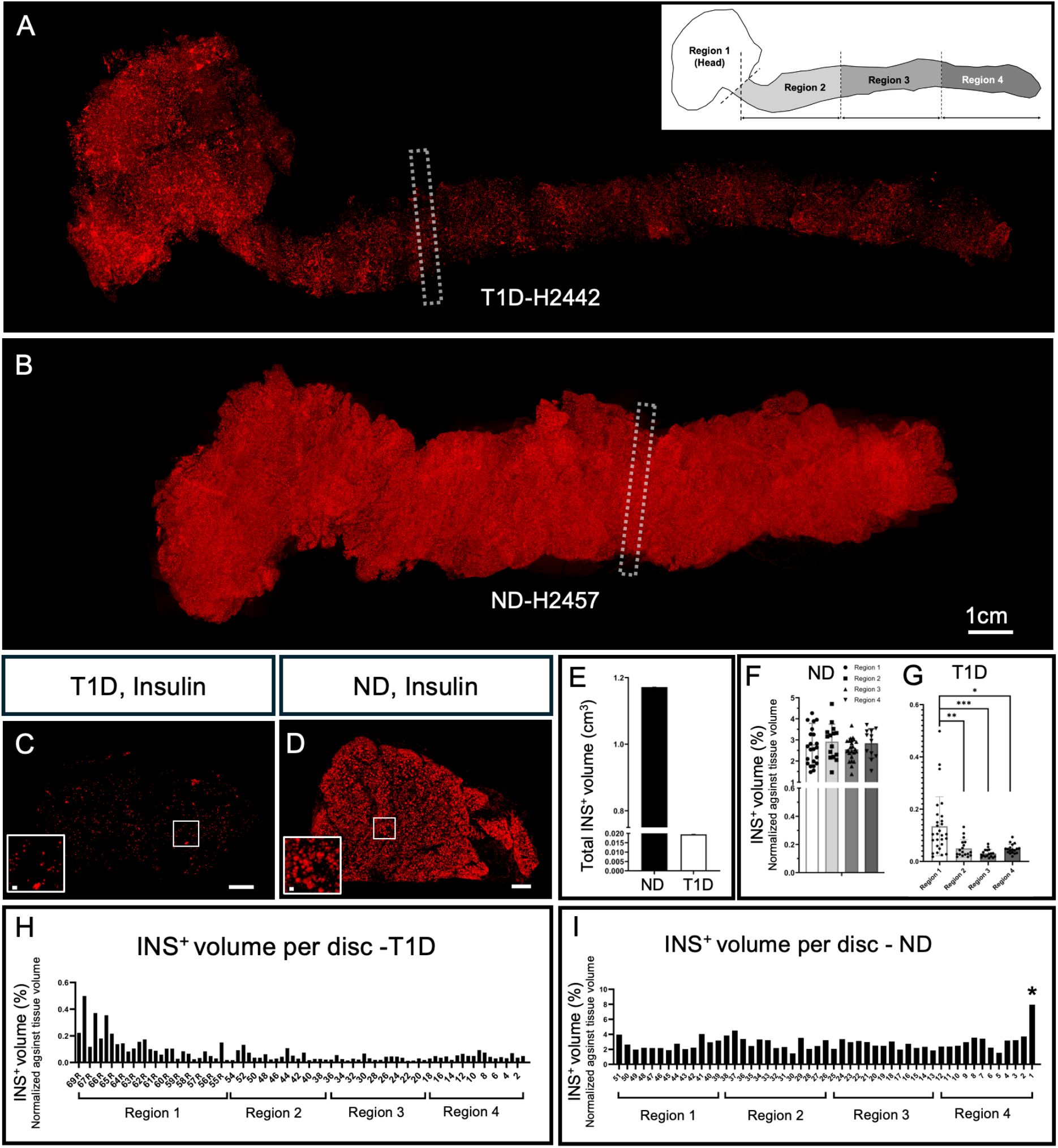
Data sets of the complete distribution of INS^+^-cells in the human pancreas from LO-T1D and non-diabetic donors. **A, B**,Pancreata from an LO-T1D (A, H2442) and a non-diabetic (B, H2457) donor were sliced into discs. The discs were individually antibody labelled for insulin (red), scanned by NIR-OPT (at 21μm isotropic resolution) and the resultant tomographic datasets aligned in 3D space, creating a fully interactive dataset (in this case a maximum intensity projection view) of the entire organ. The LO-T1D pancreas displayed an approximately 13-fold lower number of INS+ objects (ND 2.21×10^6^ vs. T1D 1.73×10^5^), and an almost 60-fold lower total pancreatic β-cell volume (ND 1.17cm^3^ vs. T1D 0.02cm^3^) compared to the ND pancreas (See [8]). **C, D**, OPT MIP images of individual discs corresponding to white boxes in (A, B). **E**, Total volume of INS^+^ cells in the depicted pancreata. Average volume of INS^+^ cells in region 1-4 as defined in inset in (A), normalized against total tissue volume. The INS^+^-cell density is 3 times higher in the T1D head region (R1) as compared to the rest of the pancreas. **F, G, H**, Relative volume of INS^+^ cells in the analysed discs (normalised against tissue volume). * In (H) represents an outlier at the end of the organ encompassing only 79 INS^+^ objects. For (F, G) a two-tailed Wilcoxon matched-pairs signed rank test ^****^ (p<0,0001) was performed. Error bars show mean ± SDE. Source data are provided as a Source Data file.

Residual β-cells appeared scattered throughout the pancreas, but with regional variations in density, particularly between the distal portion of the head and the remainder of the pancreas (**Fig. 1A, H**). Based on tomographic data segmentation, the T1D pancreas displayed approximately a 13-fold lower number of INS^+^ objects (ND 2.21×10^6^ vs. T1D 1.73×10^5^), and an nearly 60-fold lower total pancreatic β-cell volume (ND 1.17cm^3^ vs. T1D 0.02cm^3^) compared to a complete ND pancreas [8] (**Fig. 1E**). Although the ND pancreas is not aged matched with the LO-T1D sample, four other ND pancreata displayed similar BCM distribution profiles. This included a ND donor (H2522) matched for age at diagnosis of the analysed T1D donor (see **Table S1**).

The human pancreas is commonly divided into head, neck, body and tail, but the boundaries between these regions are loosely defined. To better facilitate systematic comparisons of regional features, the pancreas was divided into four regions consisting of; the head (region 1, using the indentation of the superior mesenteric vein as a boundary) and regions 2-4 consisting of portions 1/3 in length of the remainder of the pancreas (See inset **Fig. 1A**). Using this regionalisation scheme, region 1 of the LO-T1D pancreas displayed an approximately 3-fold higher β-cell density compared to regions 2-4, and even higher in the distal portion (**Fig. 1A, G, H**). In contrast, analyses of ND pancreata showed a more homogenous β-cell density across the organ head to tail (**Fig. 1B, F and I**, see also Fig. S4 in Lehrstrand et al[8]). Analyses of the β-cell density in the individual tissue discs, confirmed this picture and discs corresponding to the most distal part of the head harboured the highest percentage of β-cells normalized to the overall tissue volume (**Fig. 1H and Fig. S1**). Notwithstanding the limited sample size, this observation suggests that β-cell decay, at least in LO-T1D, can affect the organ with a high degree of regional variation with, in this case, a significantly better preservation in a region of the pancreatic head.

### Extra-islet β-cells appear less prone to β-cell destruction in LO-T1D

The applied imaging approach, allowing the entire pancreatic volume to be studied from all angles and through its entire depth greatly facilitates the identification of features that would be extremely challenging to recognize using stereological techniques, or lower resolution 3D techniques. Whereas most of the INS^+^ cells were relatively homogenously scattered in the exocrine parenchyma, local concentrations of INS^+^ cells could be observed in all regions (R1-4) of the organ on OPT images (**Fig.2A-D)**. Using these 3D data sets as a guide, regions with such clustered β-cells could be isolated, sectioned and analysed in a slide scanner. Immunofluorescence staining’s for the pan-endocrine marker Synaptophysin revealed that despite the relative distance between the INS^+^ cells within the clusters, they were not intermingled by any other endocrine islet cell-types (**Fig. 2E and 3E**), suggesting they are not part of a conventional “mixed” islet unit. The transcription factor Nkx6.1, plays a pivotal role in the specification and maturation of pancreatic β-cells, as well as in maintaining their adult functionality [13]. The INS^+^ cells of the clusters were all positive for NKX6.1 (**Fig. 2F**). Further, as determined by CD45 staining, the clusters showed no apparent signs of increased infiltration of immune cells relative to neighbouring tissue (**Fig. 2G)**, nor of increased proliferation, as indicated by MCM7 and Ki67 staining (**Fig. 2H, I** respectively).

**Figure 2.**
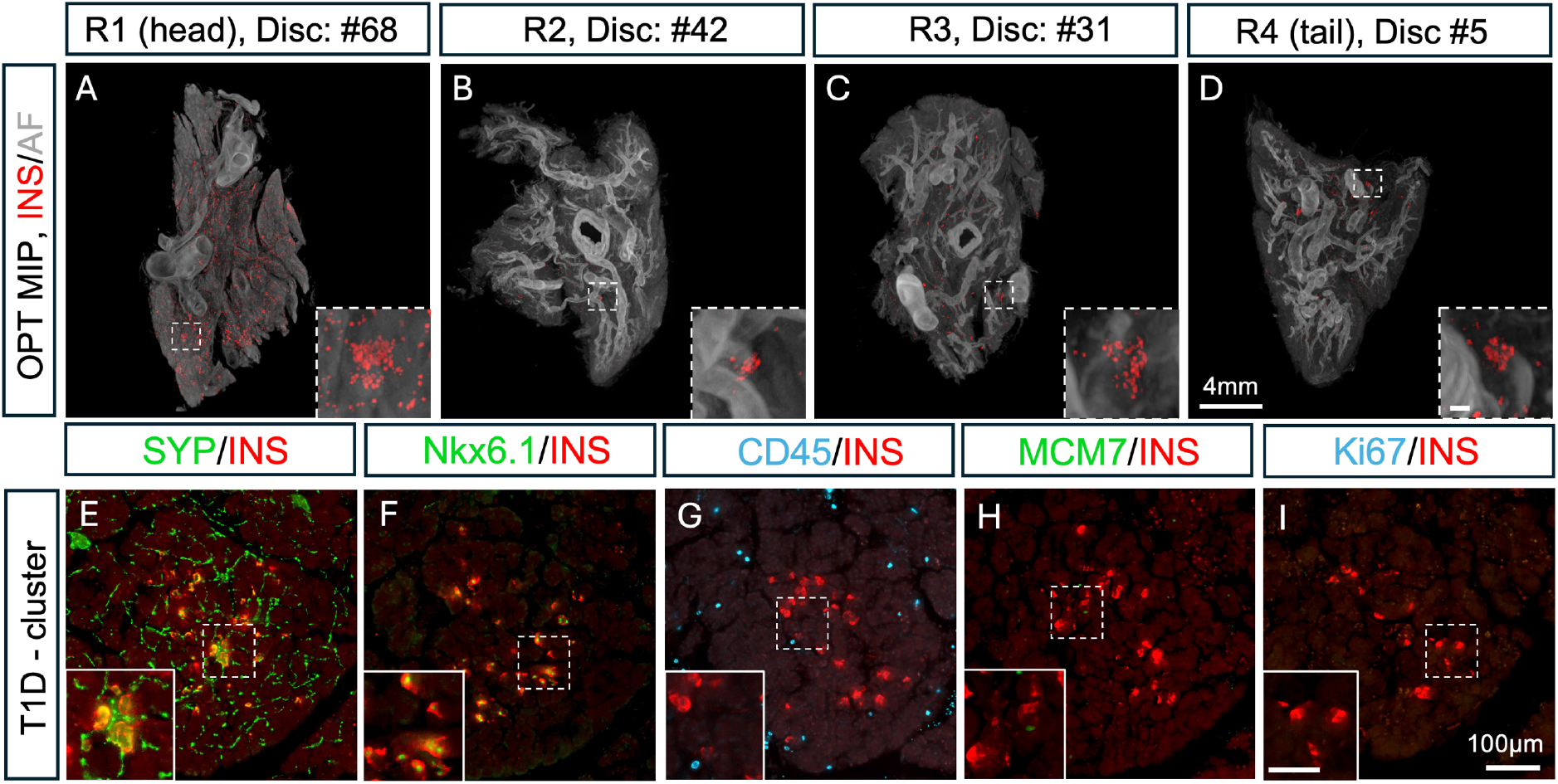
The LO-T1D pancreas harbours extra-islet β-cell clusters that are not intermingled by other endocrine cell-types. **A-D**,OPT maximum projection intensity images of representative tissue discs from region 1-4 of the pancreas shown in Fig. 1A, labelled for INS (red) in which autofluoresence (AF) from the tissue is seen in grey. Clusters of punctuated INS^+^ cells are present in all regions. **E-I**, Images from slide scanner analysis of the cluster regions showing that they are not intermingled by any other endocrine cell-types (**E**, Synaptophysin (SYP), green)., express markers of mature β-cells (**F**, Nkx6.1, green), are modestly infiltrated (**G**, CD45, blue), and that they do not show any signs of increased proliferation (**H**, MCM7, green) and (**I**, Ki67, blue). Note, green only in (E) represents innervation. Scalebar in inset in (D) corresponds to 100μm in (A-D). Scale bar in inset in (D) corresponds to 100 μm in (A-D) and scalebar in inset in (I) corresponds to 50μm in (E-I).

To quantitatively assess the spatial localisation of the residual BCM in LO-T1D in more detail, tissue discs from LO-T1D (H2422) and from ND (# H2522, matched for age at diagnosis of T1D in H2442, see **Table S1**) were labelled also for glucagon (GCG) and scanned by OPT (**Fig. 3A-D**), and smaller ROI’s by LSFM (**Fig. 3E-K**). Maximum intensity projection (MIP) views of pancreatic tissue discs from the LO-T1D pancreas showed no apparent signs of reduced numbers of GCG-expressing cells compared to discs from the ND donor, and these GCG^+^ cells were grossly organised in islet structures (Compare **Fig. 3A** and **B, E** and **F**). Segmentation of LSFM data confirmed this picture (**Fig. 3L**). By performing a classification spot analysis (see methods), clusters of INS^+^ objects (>10) residing within less than 75µm of each other could be highlighted as prominent features in the 3D data sets **Fig. S2**, for LSFM data an INS^+^ object is defined as INS^+^ cells that cannot be separated from each other at a scanning resolution of 1.6 µm). Whereas the average distance between β-cells outside the clusters was 176 μm (or 374 μm between the nine nearest neighbouring β-cells), the average distance between individual β-cells within the clusters was only 35μm, or 79μm between the 9 nearest neighbours (**Fig. S2 E, F**). These data confirm the clustered, but not directly adjacent, organisation of the β-cells observed in the 3D data. By measuring the relative volumes of INS^+^-cells I/ residing in clusters, as defined above (**Fig. 3I** and **Fig. S2**), II/ within or in direct proximity of GCG^+^ islets (**Fig. 3J**) or III/ as scattered INS^+^ cells (i.e., all INS^+^ cells not belonging to the first two categories, **Fig. 3K**, see also **Movie S2**), LSFM data showed that only 14.7% of the residual INS^+^ cell volume was associated with islet structures, whereas 85.3% was present either as clusters (22%), according to the above definition, or as scattered INS^+^ cells (63.3%) (**Fig. 3M**, white bars).

**Figure 3.**
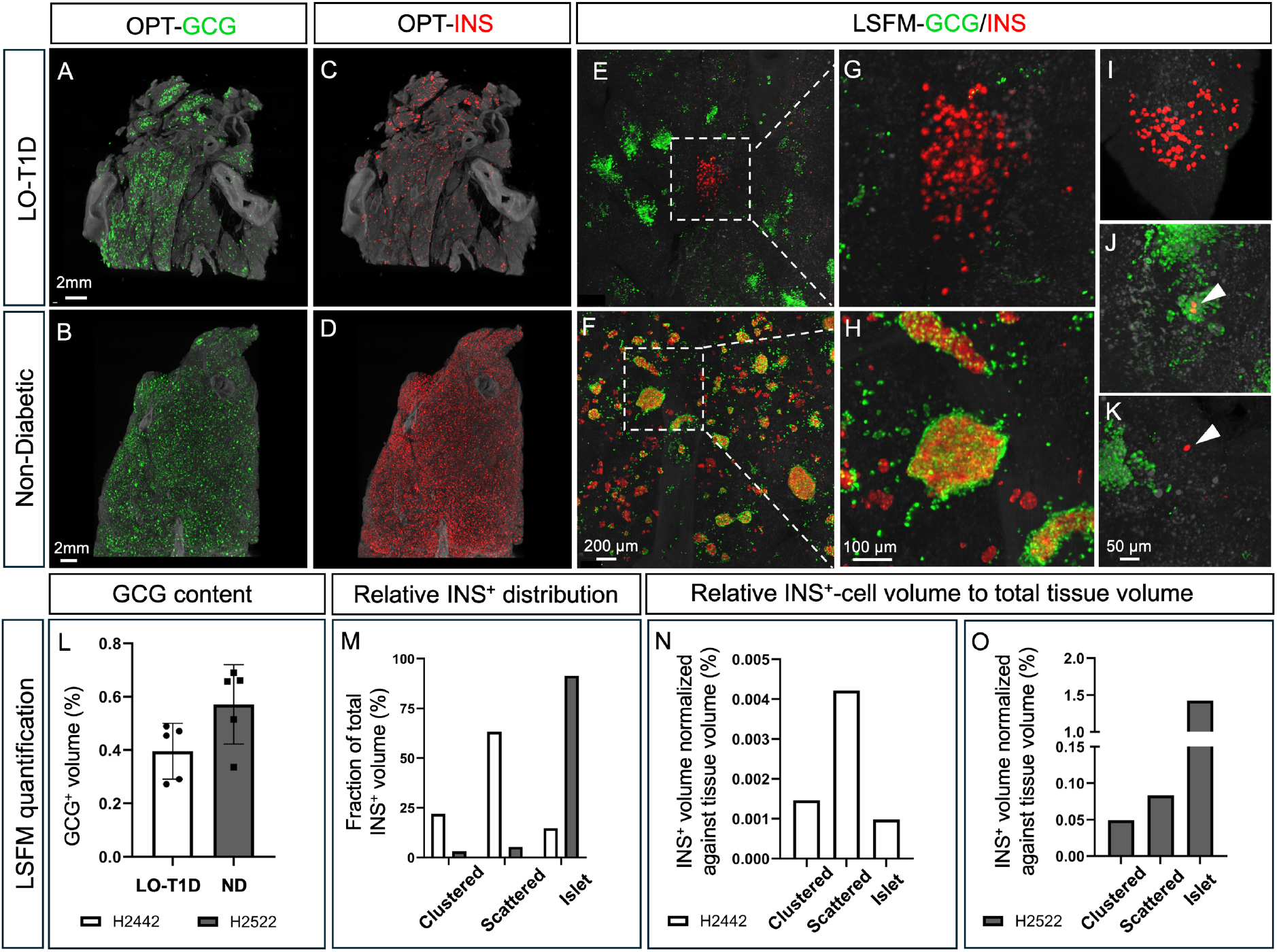
The majority of the residual BCM in LO-T1D is represented by extra-islet β-cells. **A-D**,OPT generated MIP views of representative pancreatic tissue discs (2.8mm thickness) from LO-T1D (H2442, diagnosed at 50y, disease duration 17years) (A, C) and from an ND donor (H2522, B, D), matched for age at diagnosis of H2442, stained for GCG (A, B) and INS (C, D) respectively. **E-H**, LSFM MIP views showing high resolution images of the tissues seen in (A-D). **I-K**, Illustration of the spatial location of residual INS^+^ cells in LO-T1D, which is present as clusters (I), within islets (J) or single scattered cells (J). **L**, LSFM analyses of 5 ROIs showing the volume of GCG^+^-cells normalized against total tissue volume in T1D (white) and ND (grey) bar. **M**, Bar graph showing the spatial location of the residual BCM in the pancreas in Fig. 1A (Disc #57 and #67, encompassing 10700 INS^+^ objects (H2442) and 50013 non-islet associated INS^+^ objects (H2522) respectively based on spot classification, see text for details). Only 14.7% of the residual INS^+^ cell volume is associated with islets in LO-T1D as opposed to 91.5% in the ND pancreas. **N, O**, INS^+^ volume normalized against total tissue volume for clustered, scattered or islet associated β-cells for LO-T1D (N) and ND pancreata (O), illustrating a clear shift to extra-islet β-cells in LO-T1D.

Whereas most studies of human INS^+^ cells are focused on islets, relatively little is known regarding the distribution and amount of extra-islet β-cells (i.e., individually scattered or groupings of β-cells <30μm in Ø). To assess if β-cell clusters, similar to those observed in LO-T1D, are a common feature also of the ND pancreas and to address the relative destruction of β-cells residing in the LO-T1D pancreas as clusters, scattered or islet-associated cells, we analysed an entire disc from a ND pancreas (H2522). Applying the same spot analysis as for the LO-T1D, excluding islet associated β-cells, the relative proportions of the β-cell distribution were markedly different from that of LO-T1D with 91.5% of BCM associated to islets, 5.3% as scattered β-cells and only 3.2% as β-cell clusters (**Fig. 3M**, grey bars). Next, we measured the contribution of β-cells in the above spatial categories to the overall pancreatic volume in both ND and LO-T1D pancreas. Whereas the contribution of extra-islet β-cells to the overall pancreatic volume was 0.132% (0.083 % scattered and 0.049% clustered) in the ND and 0.006% (0.004% scattered and 0.001% clustered) in LO-T1D, islet associated β-cells contributed 1.423% in the ND but only 0.001 % to the overall volume in LO-T1D (**Fig. 3 N** and **O**). Hence, comparing β-cell density between ND and LO-T1D points to a 23-fold lower extra islet β-cell density and a 1452-fold lower islet-associated β-cell density in the LO-T1D. Hence, there is a 63-fold relative change between islet associated and extra-islet β-cells, suggesting that extra-islet β-cells are much less prone to succumb than islet-associated β**-**cells comparing LO-T1D with a ND pancreas age matched with onset of diagnosis.

### Infiltration of CD45^+^ cells does not correlate to the topology of residual β-cells

To assess whether the relative variation in β-cell preservation in LO-T1D was reflected by the degree of immune infiltration, we assessed CD45 labelled discs from regions with a clear difference in β-cell density by LSFM. Immune cell infiltration was prominent and relatively uniform across the exocrine parenchyma in both “high” (disc #67R) and “low” (disc #57R) β-cell density discs (**Fig. S3**). In contrast, clear accumulations of CD45^+^ cells could be detected in the pancreatic ducts, basally along the epithelial layer (**Fig. S4**). Next, we compared the density of CD45^+^ cells in relation to the spatial distribution of residual β-cells at the single cell level. ROIs were selected to exclude vessels and ducts and were scanned at high resolution. Hereby, no significant difference could be observed in the density of CD45^+^ cells between regions with clustered β-cells or scattered β-cells only (**Fig. 4**). Hence, as judged by the density of CD45^+^-cells only, immune infiltration in LO-T1D did not display any obvious regional differences in relation to residual β-cell density and organisation, but higher concentrations could be confirmed in the ductal epithelium.

**Figure 4.**
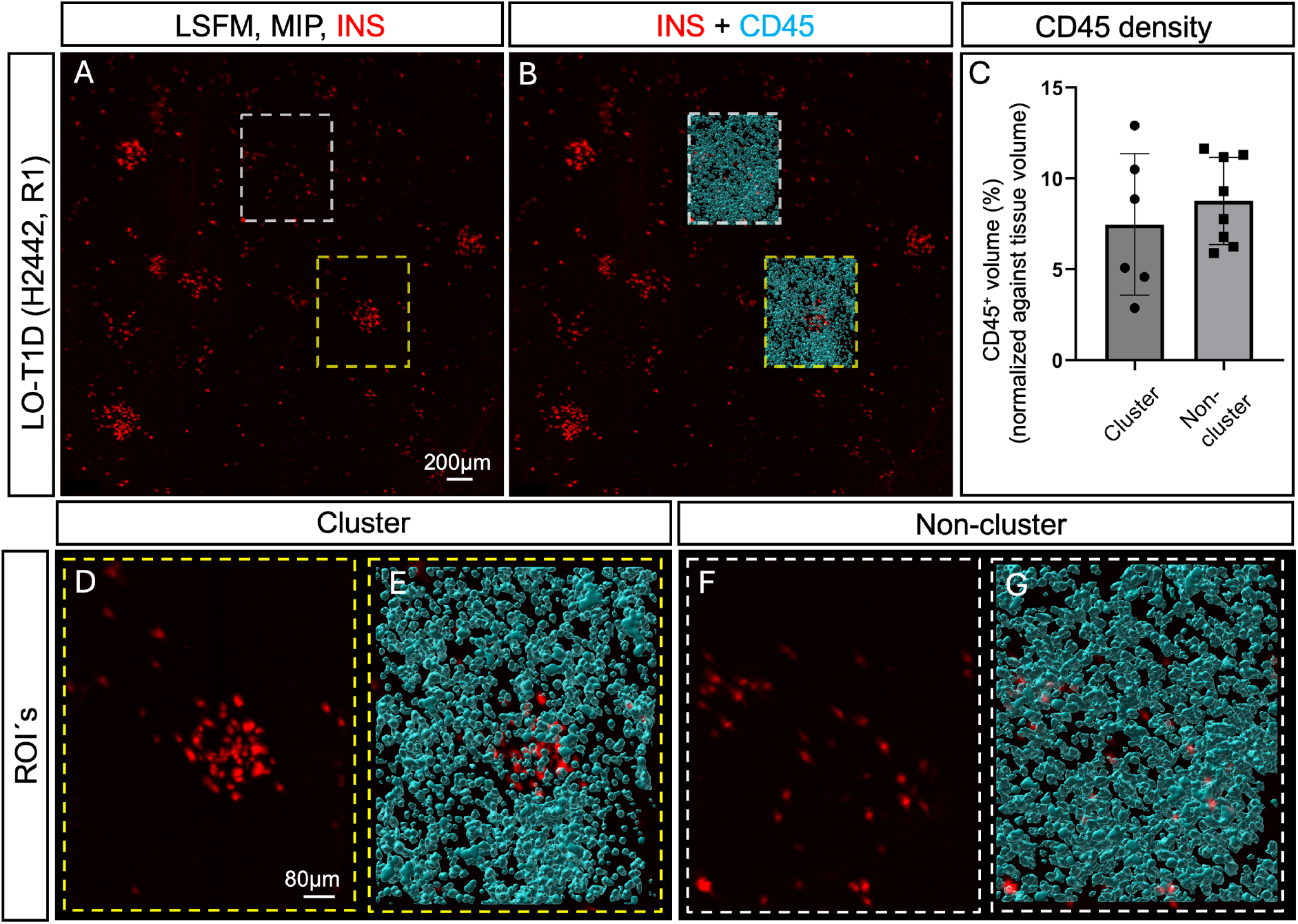
Density of immune cells appears similar in areas with clustered and non-clustered extra-islet β-cells. **A**,LSFM MIP (2.8 mm in Z) of a region of LO-T1D (H2442, Disc #67R) labelled for INS. **B**, Same image as in (A), showing examples ROIś with segmented CD45 channel (teal) covering clustered β-cells (broken yellow box) and non-clustered β-cells (broken white box). ROIś (720 x 865 x 480μm) were selected to exclude blood vessels and ducts. **C**, Bar graph showing density of CD45^+^ cells within the ROIś (CD45^+^ cell volume normalized against total tissue volume). No significant difference in CD45^+^ cell density could be observed between the two types of ROI selections. **D-G**, High magnifications of the example ROIś shown in (B). Abbreviation: ROI, Region of Interest. Scale bar in (A) is 200μm in (A and B), and scalebar in (D) is 80 μm in (D-G).

### Regional differences in loss of exocrine tissue in LO-T1D

Acinar cells decrease in number in T1D regardless of age at diabetes onset [14], and the T1D pancreas has been demonstrated to be smaller than in ND controls ([15] and refs therein). Whereas exocrine pathologies associated with T1D include acinar atrophy, fibrosis, arteriosclerosis and fat infiltration, the reason for these exocrine abnormalities in T1D and its significance for disease development is under debate [14, 16]. To address pancreatic atrophy, we calculated the tissue volume of the T1D (H2442) donor and from our previously published ND donor (H2457) using our optical tomographic datasets, which infers a close to cell level quantitative accuracy with isotropic voxel spacing. The volume of the pancreatic head (R1) constituted almost half of the total pancreatic volume in the LO-T1D donor but only 1/3 in the ND donor (**Fig. S5**).

We have previously demonstrated the possibility to use the autofluorescent (AF) properties of pancreatic tissue to visualize vascular and ductal structures in optical tomographic datasets [17, 18]. By applying this to the LO-T1D pancreas, we visualized both tissue types throughout the gland (**Fig. 5A)**. However, whereas AF intensity and the thickness of ducts and vascular walls appeared comparable to ND donor pancreas in R1 of the LO-T1D (Compare **Figs. 5B, C** with **F, G**), tomographic sections from region 2-4 displayed an apparent increase in hyperintensity of both structures (Compare **Figs. 5D, E** with **H, I**). Hence, in addition to a better regional preservation of BCM, the LO-T1D pancreatic head (R1) displayed a morphology and size better resembling the ND pancreas, whereas the remainder of the organ (R2-4) appeared more affected with regards to the above features.

**Figure 5.**
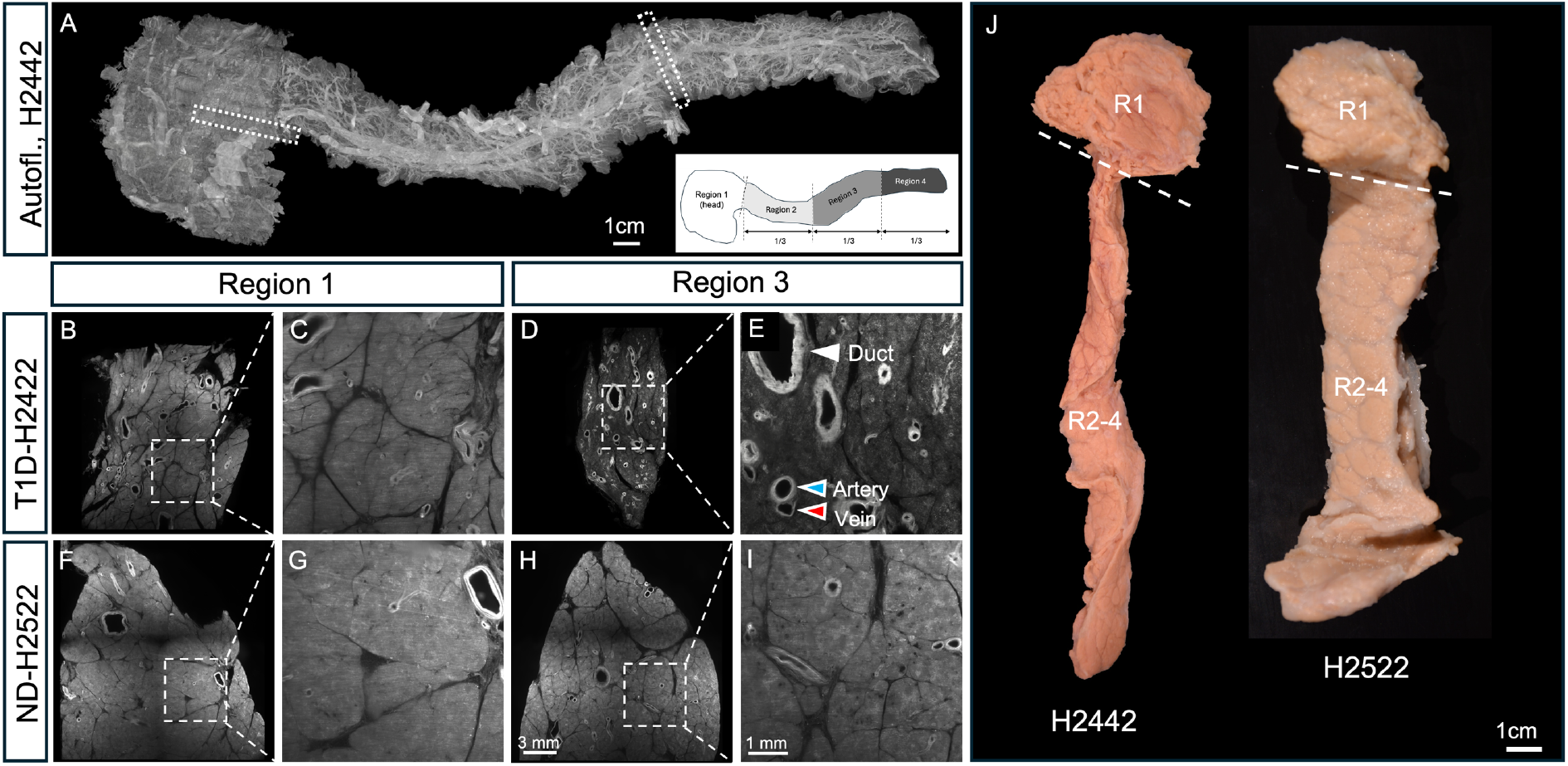
Heterogenous morphology and acinar atrophy of the LO-T1D pancreas. **A**, MIP image of the LO-T1D pancreas depicted in Fig. 1A based on the tissues autofluorescent (AF) properties. Inset illustrates the applied regionalisation scheme. Due to strong AF properties, the ductal and vascular system can be clearly visualized. **B-I**, MIP images of individual tissue discs obtained from LO-T1D (A-E) and ND (F-I) pancreas respectively. Whereas region 1 (head) display a similar morphology and AF intensity between the LO-T1D and ND pancreas, region 2-4 of the LO-T1D pancreas is clearly atrophic (see also Fig. S4) and, both vessels and ducts appear thickened and hyperintense, likely due to fibrosis of periductal and perivascular spaces. **J**, Photomicrographs of the intact pancreas displayed in (B-I) before sectioning into discs. Note the apparently slimmer appearance of region 2-4 of the LO-T1D pancreas. Scale bar in (H) is 3mm in (B, D, F, H) and scalebar in (I) is 1mm in (C, E, D, I).

## DISCUSSION

In this report, we used deep tissue optical imaging to provide a complete assessment of the residual β-cell distribution throughout the volume of a LO-T1D pancreas (H2442, see **Table S1**). Hereby, we could correlate the visual 3D data with quantitative assessments of the labelled cells throughout the organ volume. Given the scarcity of this kind of material, this provides a unique picture of disease pathophysiology on the whole organ level. Whereas we observe a dramatic reduction of β-cells, a surprisingly high number of INS^+^ objects remain (1.76×10^5^ INS^+^ objects as detected by OPT). Previous studies of LO-T1D pancreata observed a severity of β-cell loss in the order: tail>body>head [19]. Although we observe variations between analysed tissue discs, β-cell density appears relatively homogenous in region 2-4 (neck, body and tail) in our material. However, in line with the previous report [19], we see a significantly higher β-cell density in R1 (head). The tail region of the ND pancreas has been suggested to have a 2-fold higher β-cell density compared to the remainder of the pancreas [20, 21]. When employing the same deep tissue imaging approach as used here, we could in a recent study on ND donor pancreata not detect a higher β-cell density in the pancreatic tail. Instead, we found it to be relatively homogenous across the organ head to tail [8]. Notwithstanding the limited material (i.e. availability of LO-T1D tissue allowing for analysis of all regions of the pancreas), and individual variation in disease pathogenesis, our current results strengthen previous observations that β-cell destruction is spatially heterogenous in LO-T1D, and that β-cell reduction appears less prominent in the pancreatic head (R1), albeit in a regionalized manner. Noteworthy, this difference in residual β-cell density was prominent despite the apparent exocrine atrophy of region 2-4 in the LO-T1D pancreas.

Whereas individual INS^+^ cells, scattered in the exocrine parenchyma of LO-T1D and ND pancreas, have been reported previously ([6] and refs therein), our observation of distinct INS^+^ clusters, that are not intermingled by other endocrine cell types, is to the best of our knowledge not reported before. It is possible that, in previous studies, the lack of an accurate 3D imaging perspective, allowing for simultaneous detection of various endocrine cells throughout a large tissue depth, has hampered observation of these β-cell clusters. Without obvious means of lineage tracing possibilities, it is difficult to discern what these extra-islet β-cells represent. As suggested by our analyses, they are unlikely to constitute any type of reminiscent islet structure, since no other endocrine cell-types are present within or in direct vicinity of these clusters. Although they express mature β-cell markers (Nkx6.1), it is difficult to draw any conclusions on their functionality *in situ*, based on our material. Notably, although we lack information about the spatial origin of the samples within the pancreas, we could observe similar clusters by analyses of paraffin blocks from other T1D donors, (For example, see **Fig. S6**).

INS^+^ extra-islet cells have previously been shown to be reduced in T1D, although these studies suggest that this subset of β-cells are more affected in early onset T1D [6, 22, 23]. By our deep tissue imaging approach, we could determine the volume of extra-islet and islet-associated β-cells by measuring tens of thousands of INS^+^ objects. Comparing the LO-T1D pancreas (H2442) with a ND pancreas (H2522), selected to match the age of T1D diagnosis, we could observe a dramatic relative fold change of the residual β-cells in the LO-T1D coupled to their topographic anatomy. Assuming that there is no noteworthy new establishment of extra-islet β-cells (be it by transdifferentiation, proliferation or neogenesis), our data indicate that extra-islet β-cells (scattered and clustered) are roughly 60 times less prone to succumb than islet associated β-cells. Of note, the INS^+^ islet size distribution of the selected ND donor (H2522) was similar to that of ND pancreata isolated at other ages (See Fig. S4 in Lehrstrand et al[8]). Several questions regarding the nature of the extra-islet β-cells remain to be explored. Could extra-islet β-cells comprise a different, more resistant, subset of the β-cell population that have escaped the autoimmune attack?

Endogenous insulin secretion persists in long-standing T1D [24], and it has been suggested that a low-level insulin content within non-β-cells, primarily GCG^+^ cells, could be responsible for this secretion [25]. Given the relative abundance of residual extra-islet β-cells, demonstrated by our approach, we instead suggest that this population could be chiefly responsible for this secretion, or at least contribute to it. Further, since it is well established that residual β-cell function can positively affect diabetes regulation, including e.g., improved metabolic control, reduced requirement for exogenous insulin, reduced risk of late-life complications etc. Understanding why extra-islet β-cells appear better preserved in LO-T1D, and if these cells are functional, may be of key importance for a range of potential therapeutic developments. Analyses, e.g. by deep visual proteomics [26] may contribute to a better understanding of these β-cells and their possible potential for therapeutic regimens.

Reduced pancreatic volume in the range of 30-50% has been reported in LO-T1D [14, 16, 27], and our results, encompassing an entire organ with around 21μm isotropic spatial resolution, are well in line with previous observations using ultrasound, computed tomography, or magnetic resonance imaging ([27] and references therein). Notably, volume reduction has been suggested to be less severe in the pancreatic head [28-30] which was also the case in our study. Autofluorescence images of tissue discs from R2-4 in the investigated LO-T1D pancreas, exhibited dramatic differences in exocrine tissue volume compared to ND control, whereas region 1 was only marginally reduced, if at all. Further, whereas the LO-T1D R1 (head) was essentially indistinguishable from the ND control, R2-4 displayed a more stronger AF signal from ductal and vascular walls, which generally appeared thicker than in R1 of the LO-T1D and in R1-4 of ND pancreas. Although fibrosis is a common feature of both T1D and T2D, fibrosis in T1D is primarily restricted to periductal and perivascular spaces [31]. Hence, highly autofluorescent proteins, such as collagens and elastines, are likely to contribute to the hyperintense signal from these structures.

It has been demonstrated that reduction in pancreatic weight/volume. in T1D is primarily related to a reduction in exocrine cell number and not in exocrine cell size [14]. Interestingly, disease duration does not seem to stand in proportion to size reduction, which is similar regardless of time since diagnosis [14]. The mechanisms for reduced pancreas size are debatable, but insulinotropic effects on the exocrine pancreas have been suggested as a potential cause [32], and it was recently shown that insulin deficiency, independent of T1D autoimmunity, is a key factor leading to a smaller pancreas in humans [33]. In our material, β-cell loss was less severe in a part of the pancreatic head (**Fig. 1**), which has also been reported by others (see [15] and refs. therein). It has been suggested that a pancreatic polypeptide (PP) rich region of the pancreatic head exert local trophic effects that protects this region from atrophy [34]. However, the observed heterogeneity in residual β-cell distribution in our material did not appear to relate to any obvious differences in exocrine morphology within the head as judged by AF based imaging (**Fig. 1** and **5**). OPT imaging of the entire head in LO-T1D and ND pancreata, incorporating both INS and PP could potentially cast light on the relation between PP-expression, β-cell protection and pancreatic atrophy in T1D.

Further, in contrast to the ND pancreas, a high degree of immune infiltration, as marked by CD45 staining, could be detected in the exocrine tissue of the LO-T1D pancreas, and the distribution of these immune cells did not appear to have a preference for regions with higher or lower β-cell density (**Fig. S3**). Since CD8^+^, CD4+ and CD11+ cells have been reported at high levels in LO-T1D *[35]*, it is, given the promiscuous affinity of CD45 for different immune cell-types, possible that mesoscopic scale imaging incorporating these markers could refine this picture. With regards to potential insulinotropic effects, these results complicate the picture in that there is no obvious differences in exocrine morphology/size between “β-high” and “β-low” regions within the pancreatic head, despite clear differences in β-cell density.

In all, this report gives a first organ-wide view of the complete residual BCM in a LO-T1D donor with a microscopic detail. It demonstrates that the pancreas can be affected in a highly regionalized manner with regards to both β-cell mass and exocrine tissue volume. Most significantly, it suggests that BCM can be differently conserved across the length of the organ and that extra -islet β-cells may be far less prone to destruction than islet associated β-cells. Depending on their functionality, this β-cell population therefore deserves increased attention in attempts to develop new protective and/or therapeutic regimens.

## AUTHOR CONTRIBUTION STATEMENT

M. H. and J.L Contributed to study design, performed whole-mount immunohistochemical analyses, performed OPT and LSFM imaging, image and statistical analysis, and contributed to the manuscript and figures and movies. B.M performed image and statistical analyses. T. A. contributed to the manuscript, study design and immunohistochemical analyses. O. K. Initiated the study, contributed to the study design and commented on the paper. U. A. Initiated and supervised the study and wrote the paper.

## ACKNOWLEDGEMENTS

The authors thank Mrs. S. Ingvast, Drs. C. Nord and W.I.L. Davies for technical assistance and Dr. S.M.A. Willekens for helpful comments on the manuscript. This work was supported by the Kempe Foundations (JCSMK24-0063 to U.A.), the Swedish Research Council (2023-02221 to O.K), Umeå University Biotech Grants (FS 2.1.6-2026-20 to U.A.), a Umeå University Strategic Research Grant (FS 2.1.6-74-24 to U.A.), the Swedish Childhood Diabetes Foundation/Barndiabetesfonden (to U.A. and O.K.), the Novo Nordisk Foundation (NNF21OC0069771, 0084520, NNF24OC0092100 to U.A.), the Diabetes Wellness Foundation Sweden (PG21-6566 to U.A.), the Ernfors Family Fund (2023 to O.K.), Nils Eric Holmstens forskningsstiftelse (2023 to O.K.) and Diabetesfonden (DIA2024-914 to UA).

## Notes

### Competing Interest Statement

The authors have declared no competing interest.

