## Supplementary Data for "Deep tissue optical 3D imaging reveals preferential preservation of extra-islet β-cells in late-onset Type 1 Diabetes"

Joakim Lehrstrand *et al.*

### The PDF file includes:

Material and Methods
Figures S1 to S6
Table S1
Description of Additional Supplementary Files

### MATERIALS AND METHODS

#### Ethics declaration

All work involving human tissue was conducted in accordance with the Declaration of Helsinki [36] and in the European Council's Convention on Human Rights and Biomedicine [37]. Consent for organ donation for use in research was obtained from the donor prior to death via the Swedish National Donor Registry (https://www.socialstyrelsen.se/en/apply-and-register/join-the-swedish-national-donor-register/) or from relatives of the deceased donors conferred by the donor's physician and documented in their medical records. The study was approved by the Regional Ethics Committee in Uppsala, Sweden (Dnr 2017/1471-32, 2023-01845-01).

#### Pancreas isolation and preparation.

The T1D and ND pancreas (see **Table S1** for clinical information and ischemia times) were obtained as donations post-mortem from the Nordic Network for Clinical Islet Transplantation (NNCIT). Shortly after the donor's death, the fresh pancreata were separated from the duodenum in Ringer's acetate (Braun Melsungen AG Hessen, Germany). Before immersion fixation in 4% formaldehyde (Solveco, Roserberga, Sweden) for 24h, the organs were washed rigorously in 1x PBS (Medicago AB, Uppsala, Sweden). The fixative was replaced with fresh 4% formaldehyde and the glands were incubated for another 24h, followed by stepwise dehydration into ethanol (2x75% v/v and 2x96% v/v, VWR chemicals) at 4°C. At this point, the dehydrated organs were transported at room temperature and extensively washed in 96% (v/v) ethanol at 4°C upon arrival. The organs were repeatedly washed in fresh 96% (v/v) ethanol on an orbital shaker at 4°C until residual fat was removed and the ethanol stayed transparent. Thereupon, images of the intact organs (**Figs. S1** and **S5**) were taken with a Nikon D5200 camera.

#### Tissue processing for optical 3D imaging.

A custom designed slicing matrix (slicing thickness 2.8mm) was generated in Tinkercad (Autodesk, USA) [8]. The matrix was 3D printed with a 1.75mm thick PLA filament (Prusa i3 MK3S, Prusa Research, Czech Republic). Before mounting the entire organs in 37°C warm 1.5% low melting point agarose (LMA) (Lonza™ SeaPlaque™ Agarose, Lonza, USA) the head (region 1, see inset in **Fig. 1A**) of the pancreas was separated to allow fitting into the slicing matrix. Region 1 and regions 2-4 were separately mounted and cured at 4°C for 4-6h. Once the agarose was solidified, the pancreas was sliced into discs, followed by removal of the surrounding agarose. An image was captured of each disc as reference for downstream 3D stitching (**Fig. S1**). The discs from region 1 of the LO-T1D pancreas were divided in two to fit within the field of view (FOV) of the NIR-OPT scanner at pre-determined magnification. Each pancreatic disk was subsequently washed at least three times for 2h in 100% Methanol (cat: 67-89-4, Fisher Scientific, Sweden) on slight rotation to further remove excessive lipids and remaining agarose. The samples were then subjected to 5 freeze/thaw cycles (brought to -80°C for 1h and back to room temperature (RT) in each step) in 100% methanol for permeabilization, followed by ON incubation in a bleaching solution (30% stock solution of $H_2O_2$; Cat. No. H1009; Sigma-Aldrich, Merck, Germany) at a final concentration of 15% (v/v), 16,7% (v/v) dimethyl sulfoxide (DMSO; Cat. No. D5879; Sigma-Aldrich, Merck, Germany) in MeOH (3:1:2 ratio). After this incubation, the samples were again incubated in bleaching solution for another 6-8h, followed by 2 washes in 100% methanol for 2h each at RT.

For paraffin embedded tissues (**Fig. S6**), the samples were deparaffinized as described in [38]. In Brief, the samples were subjected to a minimum of two 1h xylene treatments at 37-50°C or until all paraffin had been solubilized, followed by a 1h xylene incubation at RT and removal of residual paraffin was performed in a downgraded series of ethanol solutions into 1x PBS (33% 1xPBS, 66% PBS and finally 100% 1xPBS), 1h per step.

**Whole mount immunohistochemistry**

Whole mount immunolabelling was performed essentially as previously described [17]. In short, pancreatic slabs were stepwise (33%, 66%, 100% and again 100%, 1h each step) rehydrated from methanol to TBST (0.15M NaCl, 0.05M Tris-HCl pH 7.5, 0.1% Triton® X-100 (cat: 108603, Merckmillipore, Germany)), NaCl and Tris-HCl produced in-house (TBK, Norrlands Universitetssjukhuset, Umeå)) on rotation. The rehydrated pancreatic discs were next incubated in blocking solution (TBST, 10% heat-inactivated goat serum (cat: CL1200-500, Cedarlane, Canada), 5% DMSO (cat: D5879, Sigma-Aldrich, Merck KGaA, Germany) and 0.01% sodium azide) for 2 days at 37°C with slight orbital shaking. Next, the samples were incubated with primary antibodies, including guinea pig anti-Insulin/Pro-Insulin (cat: 16049, PROGEN Biotechnik GmbH,

Germany, diluted 1:3000) and for select discs; rabbit anit-glucagon (cat. no.: HPA036761, Atlas Antibodies, Stockholm, Sweden, diluted 1:1000) in blocking solution at 37°C for 7 days on slight orbital shaking, and thereafter washed in TBST, 5 times 1h on a rotator at RT. Secondary antibody incubation was performed similarly as the primary using donkey anti-guinea pig IRDye 680 (cat: 926-68071, Li-Cor, USA, diluted 1:250) and goat anti-rabbit Alexa Fluor™ 594 Invitrogen™ (cat: A11012, Thermofisher, USA, diluted 1:500). To remove possible fluorophore precipitates, the secondary antibody solutions (blocking and secondary antibody) were first filtered through a 25mm wide Acrodisc® with a pore size of 0.45µm (cat: 4614, Pall Corporation, USA). The samples were then washed again rigorously 5x1h in TBST at RT on rotation. Next, any residues (like occasional fibers) were removed prior to washing in distilled water and mounting in 1.5% LMA at 37 °C to generate a scaffold for the sample holder in NIR-OPT. The agarose was then allowed to solidify for 3-4h at 4 °C. Excessive agarose was later trimmed to fit the NIR-OPT sample holder and samples were dehydrated by 5 washes of 100% methanol for 1-3h each step on rotation. Samples were optically cleared using BABB, a 1:2 mixture of benzyl alcohol (cat: 100-51-6, Merck, Germany) and benzyl benzoate (cat: 105860010, Acros organics, USA) by replacing BABB 5 times over 2.5 days before optical 3D imaging.

**Near infrared Optical Projection Tomography (NIR-OPT) and Light Sheet Fluorescence Microscopy (LSFM)**

3D image acquisition of the agarose embedded pancreatic disks were effectively performed in an in-house build NIR-OPT scanner [9] as previously described [8, 17]. In brief, the optical tomograph was built from a Leica MZFLIII stereomicroscope (Leica, Wetzlar, Germany) with a tilted mirror and step motor for sample rotation with an Andor iKon-M camera (Andor technology, United Kingdom) and attached to a CoolLED pE-4000 fluorescence light source (Ludesco Microscoped, USA). The scans were performed at a magnification of 1.25x, rendering a final isotropic voxel size of 21 µm. Filter sets include Ex: 425/60 nm, Em: 480nm LP for autofluorescent (AF) signal, Ex: 565/30nm, Em: 620/60nm for glucagon and Ex: HQ 665/45nm, Em: HQ 725/50nm for insulin. Exposure times were 800ms for the AF, 6000ms for glucagon and 8000ms for the insulin channel.

Subsequent high resolution ROI analysis of previously NIR-OPT scanned samples was performed using a commercially available LSFM: the UltraMicroscope II

(Miltenyi Biotec, Germany), which included a 1x Olympus objective (Olympus PLAPO 2XC) with a lens corrected dipping cap MVPLAPO 2x DC DBE attached to an Olympus MVX10 zoom body and 3000 step chromatic correction motor. Consistent ROIs of T1D and ND samples with the entire z-axis depth were captured with a zoom factor of 0.63x, optical sectioning step-size of 10 µm, numerical aperture of 0.14, and a dynamic focus across the field of view of 10 images per section plane. Light sheets were merged using the built-in projection function. Exposure times for all channels was kept at 300ms with filter sets for channels AF594 (CD45 **Fig. S3**) Ex: 580/25, Em: 625/30 and for AF (**Fig. 5** and **Fig. S3**) Ex: 470/40, Em: 525/50, respectively. Optical section images were generated in industrial *.ome.tif format by the ImspectorPro software (version 7.0.124.9 LaVision Biotec GmbH, Germany). Finally, the resultant LSFM data sets were directly converted to *.ims files using the Imaris file converter (v10.2, Bitplane, UK). Imaging of insulin and glucagon distribution (**Fig. 3** and **Figs. S2**, **S4** and **S6**) was performed in the BLAZE (Miltenyi, Biotec, Germany) using an 4x objective lens NA 0.35 MI PLAN. Images were captured at 1.0x magnification using filter combination Ex. 710/75 Em. 810/90 for insulin, Ex. 560/40 Em. 620/60 for glucagon and Ex. 470/30 Em. 525/60 for anatomy at a sheet NA of 0,061 with step size of 6µm. Tiled images were stitched using MACS iQ View 3D (ver. 1.2, Miltenyi, Biotec, Germany).

**3D image processing and reconstruction for NIR-OPT data**

Post-NIR-OPT processing was performed essentially as described [8, 17]. In brief, the acquired OPT image data were processed using the “DSP-OPT” software package (https://github.com/ARDISDataset/DSPOPT), which includes image processing assisted algorithms specifically for OPT [17]. Beforehand, the pixel range of OPT frontal image projections was cut for each pancreatic disc to display minima and maxima. Next, a contrast limited adaptive histogram equalization (CLAHE) algorithm with a tile size of 48 by 48 with sharpening for the insulin and glucagon channel and A-value tuning (alignment of axis of rotation using a Discrete Fourier Transform Alignment algorithm) for all channels was implemented from the DSP-OPT software package, aiming to improve the signal to noise ratio and 3D image alignment. Consequently, the processed frontal projections were reconstructed into tomographic image sections using NRecon software (v1.7.0.4, Skyscan Bruker microCT, Belgium), which included additional misalignment compensation at the axis of rotation and ring artifact reduction. Finally, reconstructed image data sets were converted to Imaris files for quantification and analysis.

In Imaris 3D analysis software, each pancreatic slab was 3D cropped and re-oriented to fit to its original location post-matrix sectioning (**Fig. S1**). Quantification in Imaris of anatomy (AF) and insulin positive voxels was performed using an automated batch processing pipeline with background subtraction and the provided surfacing method from Imaris. Hereby, $INS^+$ objects were included with a voxel threshold value >5 and objects below a volume of 3000 $\mu m^3$ were excluded to remove noise. In addition, artefacts, such as fibers outside the anatomy channel, were also manually excluded.

**3D stitching the entire pancreas volume**

The 3D data sets of the non-overlapping pancreas slabs were oriented in Imaris to fit the post-matrix slicing images. Manual 3D stitching was performed in Imaris by adding one pancreatic disk at a time. By alignment of individual ducts and vessels in the AF channel of each slab the entire T1D pancreas (82 slabs with a total of 164 channels) was reconstructed in 3D space.

**Classification and distance measurements of extra-islet b-cells**

High resolution LSFM scans of ROIs were analysed in Imaris (see **Fig. 3E-K**). Volumetric surfacing of $GCG^+$ signal and density quantification was performed in Imaris as previously described [8]. To describe the distribution of extra-islet β-cells, isolated "punctuated" $INS^+$ clusters (**Fig. 3G**) were arbitrarily classified using the spot segmentation function in Imaris. As such, $INS^+$ objects with ≤75 μm to the nearest $INS^+$ object with 5 and 9 nearest neighbour set to 120 and 100 respectively, containing more than 10 objects, were classified as "clustered" cells. Further, any $INS^+$ spots associating (≤45 μm distance) within glucagon volume were classified as "islet" cells. Finally, objects outside these grouping parameters were considered "scattered" (See **Fig 3I-K** and **Fig. S2**). From this spot classification, the segmented $INS^+$ volume could be sorted into the three categories by using the Imaris overlap function. This allowed quantification of distance parameters between $INS^+$ objects, such as distance to nearest $INS^+$ neighbour or average distance to the nine nearest $INS^+$ neighbours (see

**Fig. S2 E** and **F** respectively), volume fractions and density (**Fig. 3M-O**) for each subcategory. Data was extracted as *.xml files for statistical analysis.

#### Quantification of CD45+ immune infiltration

To evaluate the density of $CD45^+$cells we evaluated ROI´s of720 x 865 x 480 μm (xyz) by LSFM. Size was set to include the largest cluster. Spot analysis to define cluster and non-clustered β-cells was conducted as described above. ROIs were placed with clusters centred; non-clustered β-cell ROIs was preferentially selected in proximity to cluster ROIs. All ROIs were chosen to exclude areas with major ducts or vessels. Volumetric rendering of signal was performed in Imaris with minimum voxel threshold set to >10 to remove noise. Volumetric data was extracted as *.xml for statistical analysis.

#### Post 3D imaging histology

Pancreatic discs with extra-islet b-cell clusters identified in OPT, were washed several times in 100% MeOH to remove all traces of BABB, followed by stepwise rehydration in a series of 70%, 50%, 30% and 10% (v/v) ethanol to 1 × PBS for 1h at RT with gentle shaking per step. Agarose was removed by first washing the slabs in 0.29 M sucrose (Cat. No. 10319003; Fisher Scientific, Sweden) in 1 × PBS at 57 °C, then two further washes with 0.29 M sucrose solution at RT with careful manual removal of agarose where required. After agarose removal, tissues were cryoprotected by incubating in 30% (w/v) sucrose in 1 × PBS overnight at 4 °C, embedded and snap frozen in NEG-50 (Cat. No. 11912365; Fisher Scientific, Sweden) and stored at −80 °C. 20um thick serial sections were collected onto SuperFrost Plus glass slides (Cat. no. 10149870; Fisher Scientific, Sweden), air dried at RT and washed in TBST for 10 min. Tissue sections were blocked in 10% fetal bovine serum (FBS; Cat. no. 11550356; Sigma-Aldrich, Merck, Germany) for 1h at RT and re-stained with insulin (diluted 1:10000) combined with either Rabbit anti -synaptophysin (Cat. no. ab32127, Abcam, UK, diluted: 1/2000), -Nkx6.1 (Cat. no. NBP1-49672, Nous biotechne, US, diluted: 1/500), -CD45 (Cat. no. A700-012 Fortis life sciences, US, diluted: 1/2000), -MCM7 (Cat. no. HPa003898, Atlas Antibodies, SWE, diluted:1/500) or -Ki67 (Cat. no. 9129S, Cell signaling technology, US, diluted 1/500) (**Fig. 2**), in blocking solution at RT overnight. Slides were washed 3 × 5 min in TBST and incubated with 4′,6-diamidino-2-

phenylindole (DAPI) and the following secondary antibodies in blocking solution for 2h at RT: Alexa Fluor 488® goat anti-guinea pig IgG H&L (Cat. No. ab150185; Abcam, UK; diluted 1:500 and Alexa Fluor 594® donkey anti-rabbit IgG H&L (Cat. No. 711-585-152; Jackson ImmunoResearch, UK; diluted 1:500). After incubation, slides were washed in 3 × 5 min in TBST and mounted with Vectashield® mounting medium (Cat. No. H-1000; Vector Laboratories, USA).

All 2D sections were scanned using the automated Axio Scan.Z1 Slide Scanner (ZEISS, Germany), equipped with a Colibri 5/7 light source. DAPI, AF488 and AF594 were imaged using an Axiocam 506 microscope camera (ZEISS, Germany) with a Plan-Apochromat 20×/0.8 M27 objective and the following filters for: DAPI; 90 HE DAPI (Ex 353, Em 465), AF488; 90 HE GFP (Ex 493, Em 517), and AF594; 64 HE mPlum (Ex 590, Em 618). For each section, 15 Z-slices were taken at a range of 10 µm, with scans being saved in *.czi format and analyzed using ZEN (blue edition) microscopy software (version 3.7.97; ZEISS, Germany). ROIs containing INS+ clustering cells (**Fig. 2**) were selected in ZEN and Z-stack images presented as orthogonal projections (maximum) with deblurring (strength, 0.5; BlurRadius, 15; and sharpness, 0).

**Statistical analyses**

All numerical data from individual samples from OPT scans ($INS^+$ volumes and anatomy volume) and LSFM scans (distance measurements of $INS^+$ objects) were extracted from Imaris as *.csv files. By use of the query import function, multiple parameters (e.g., volume, area or diameter) were imported and organized in Excel (Microsoft, office 365, version 2505) with an individual ID, pancreatic region ID and pancreas ID. For regional density comparisons (see **Fig. 1G and H**) the average of each pancreatic slab was employed, thus each dot in the graphs represents multiple thousand objects.

In general, graphs, distribution curves and significant tests were generated and performed in GraphPad Prism (v10.5.0, LCC). For **Fig. 1G** a Shapiro–Wilk test for normality was applied for each column, as the test for normality (gaussian distribution) was not passed, we applied a Wilcoxon matched-pairs signed rank test for comparison. In **Fig S2** a two-tailed paired t-test was applied. *P*-values are reported as: $P > 0.05$ (ns), $P \leq 0.05$ (*), $P \leq 0.01$ (**), $P \leq 0.001$ (***), $P \leq 0.0001$ (****).

### SUPPLEMENTARY FIGURES

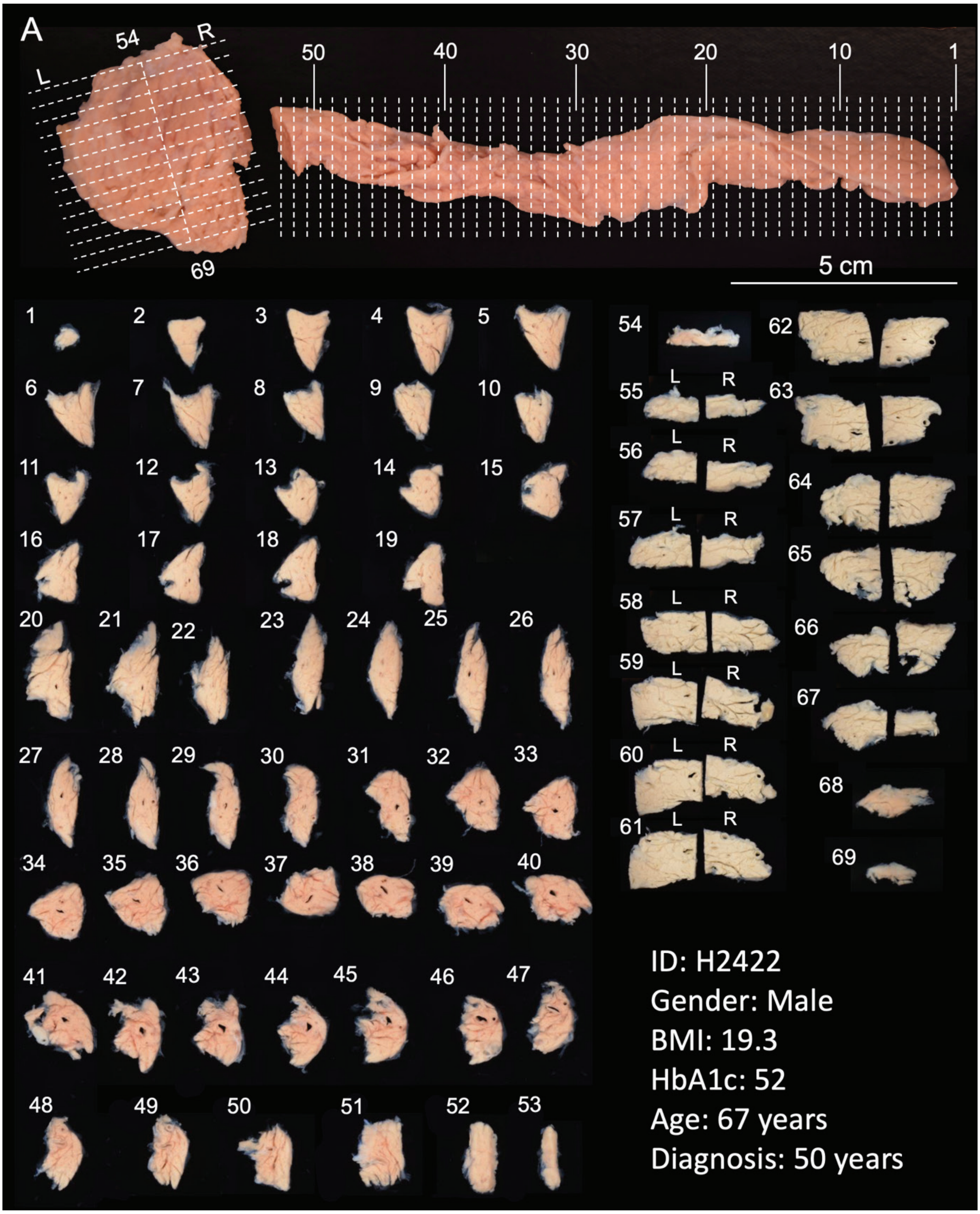


**Figure S1. Slicing and documentation of pancreatic discs for labelling and 3D analysis**. **A,** Human pancreas (H2442) was sliced in a 3D-printed matrix (grid size 2.8 mm). **B,** Photomicrographs of individual discs (1-69) from tail (1) to head (69). Note, sections 55-67 were cut in half to facilitate NIR-OPT scanning. Abbreviations, L: left, R: right.

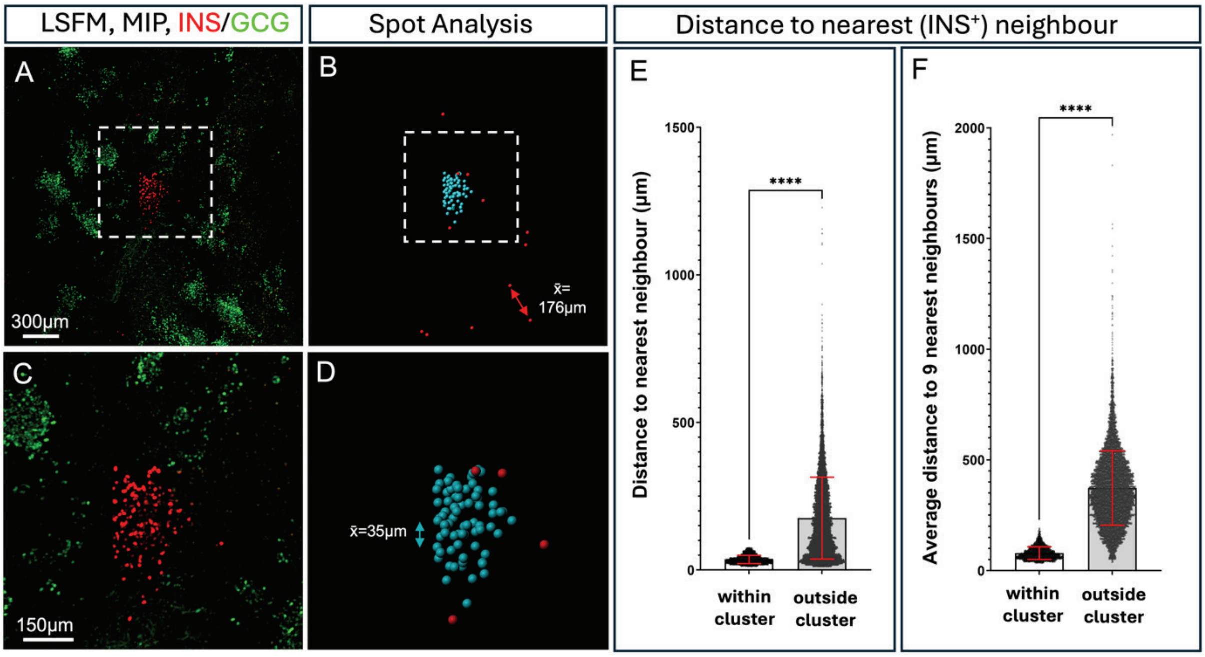


**Figure S2. Identification and 3D distance quantification of β-cell clusters in LSFM scanned LO-T1D pancreas. A,** Representative MIP 3D view of a high resolution LSFM scan of a region of interest (ROI) from R4 of the pancreas displayed in Fig. 1, showing INS$^+$ volumes (red) and GCG$^+$ volumes (green). Image in (C) corresponds to the region indicated by broken white line in (A). Isolated, yet clustered, β-cells appear in close distance to each other. **B,** Image of spot distance categorisation based on INS$^+$ voxels performed in Imaris. By filtering segmented INS$^+$ spots into groups of ≤75 µm distance of each other, isolated β-cell clusters (pseudo coloured in teal) with ≥10 objects per group are displayed. Ungrouped INS$^+$ spots and groups with less than 10 objects are displayed as red spots. Average distance to the nearest INS$^+$ neighbour within the β-cell clusters was ~35µm and average distance between ungrouped INS$^+$ objects was ~176 µm. **F**, **G,** Bar graph showing the average distance to the nearest neighbouring INS$^+$ object (F) and to the nine nearest INS$^+$ objects (G) for clustered and non-clustered INS$^+$ objects respectively. Error bars represent SD. **** $p<0.0001$.

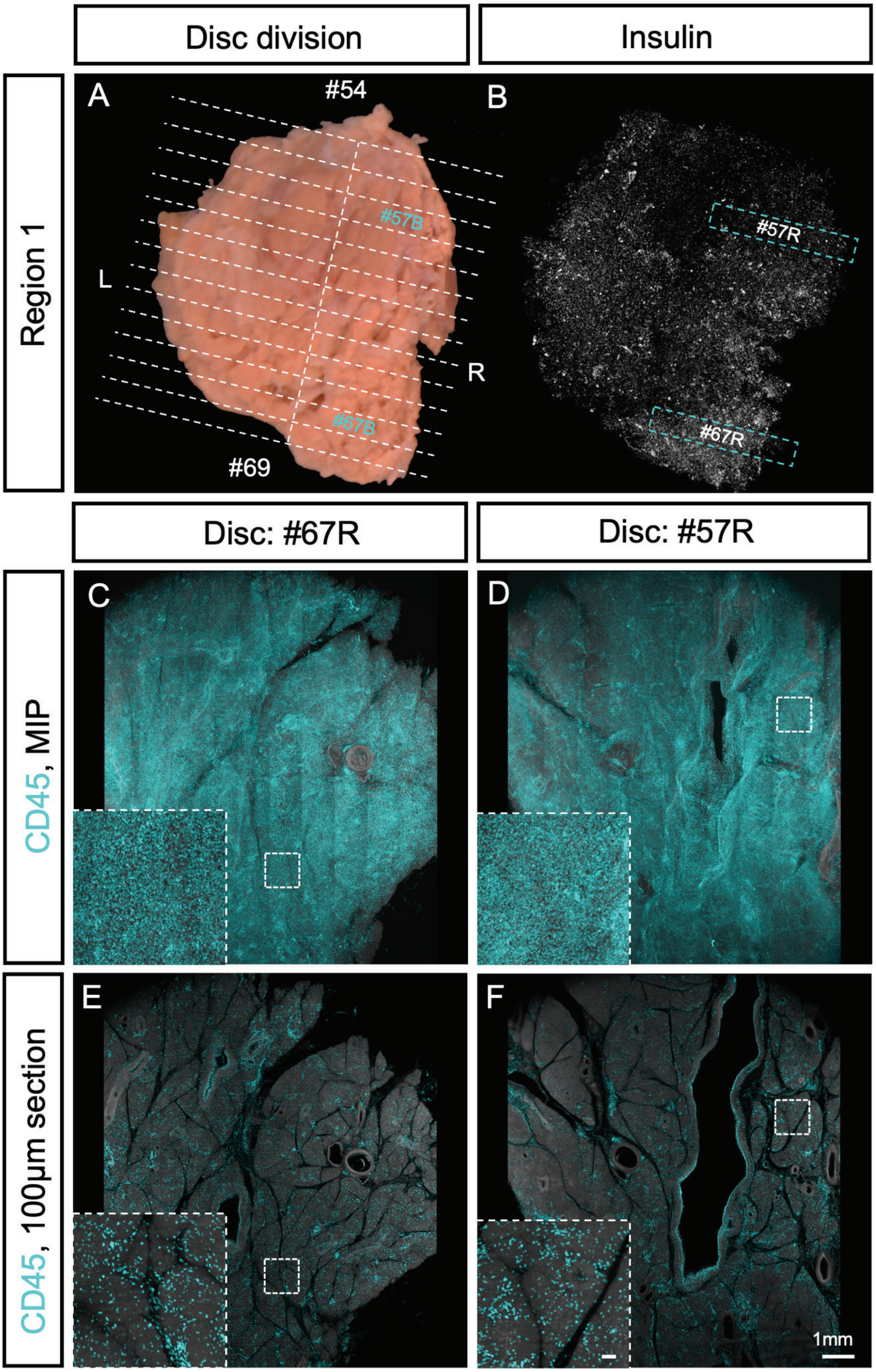

Disc division
Insulin
Region 1
A
#54
#57B
L
R
#67B
#69
B
#57R
#67R
Disc: #67R
Disc: #57R
CD45, MIP
C
D
CD45, 100µm section
E
F
1mm

**Figure S3. Immune-infiltration appears homogenous in areas of high and low β-cell density. A**, Photomicrograph of region 1 (pancreatic head) from LO-T1D donor H2442 illustrating the spatial origin of the analysed discs. **B**, OPT MIP image showing insulin staining (white) of all the discs in (A) aligned in 3D space. Teal colour marks area with higher (#67R) and lower (#57R) β-cell density (See also Fig. 1A). **C, D**, LSFM MIP image of disc #67R (C) and #57R (D) stained for CD45 (thickness 2.8mm). **E, F,** 100μm optical LSFM sections corresponding to a (C, D). Insets in (C-F) correlates to the regions indicated by broken white line. There is no apparent difference in CD45 staining between low and high β-cell density regions. Scale bar in inset in (F) corresponds to 100μm in (C-F).

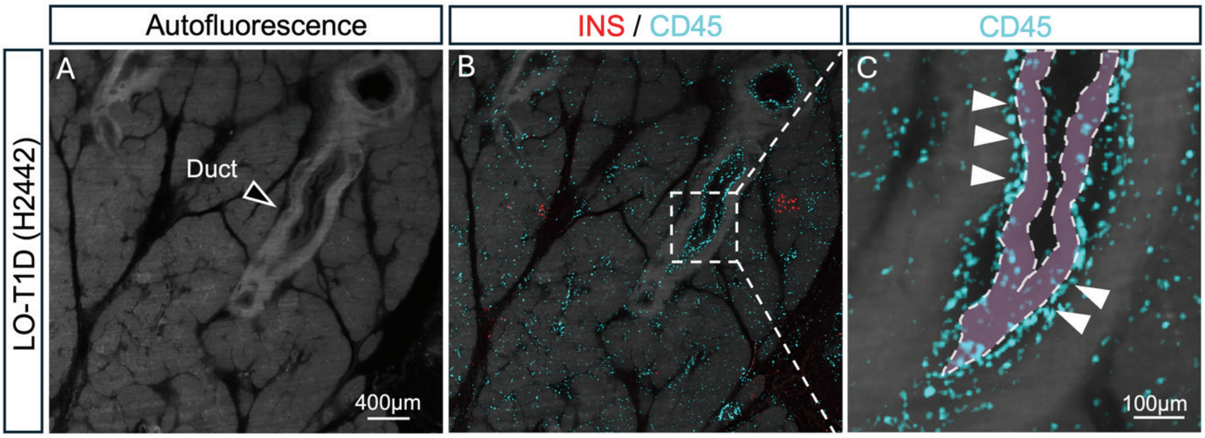


**Figure S4. Immune cells are infiltrating the inner linings of the pancreatic ducts.** **A**, LSFM section (10µm), showing the AF signal from LO-T1D (H2442). Closed arrowhead is pointing to the hyperintense signal from a pancreatic duct. **B**, Same region as in (A), labelled for INS (red) and CD45 (teal). **C**, Image corresponding to the broken line box in (B). The inner lining of the pancreatic duct is highlighted in (transparent purple). White arrowheads are pointing to immune cells that has infiltrated the inner lining of the duct.

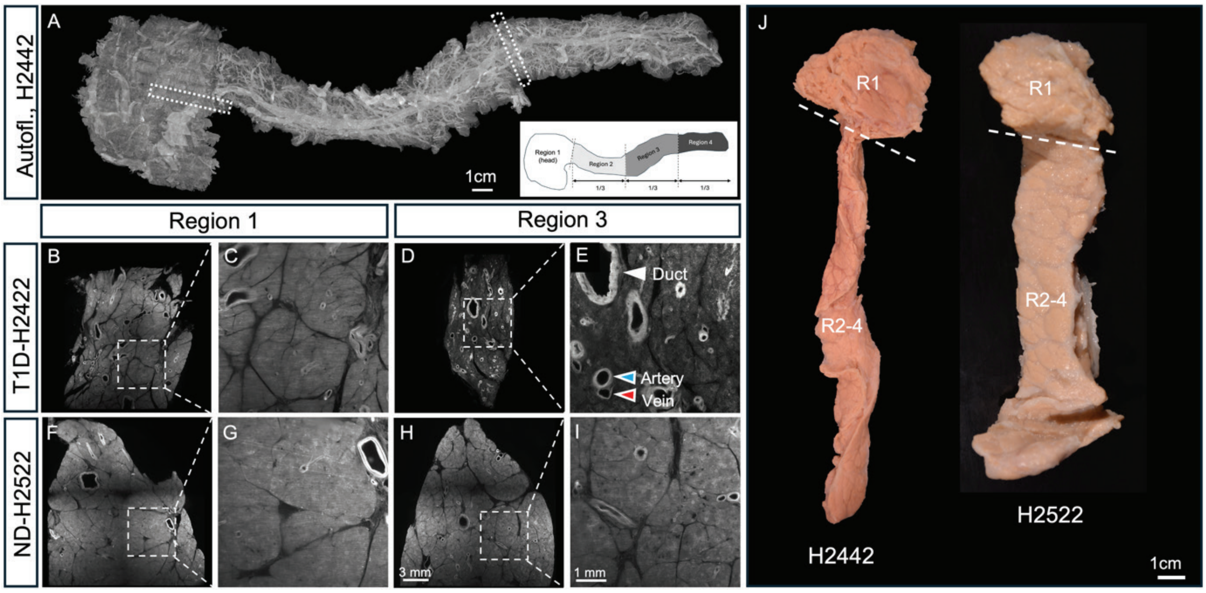


**Figure S5. Pancreatic atrophy is predominantly a feature of the pancreatic body and tail in LO-T1D. A,** Bar graph displaying pancreatic tissue volume of region 1 and regions 2-4, respectively, in a ND (H2457 - black) and LO-T1D (H2442 - white) donor pancreas measured by OPT imaging of the entire organ (See Table S1). **B**, Photomicrographs of the pancreata that were analysed in (B).

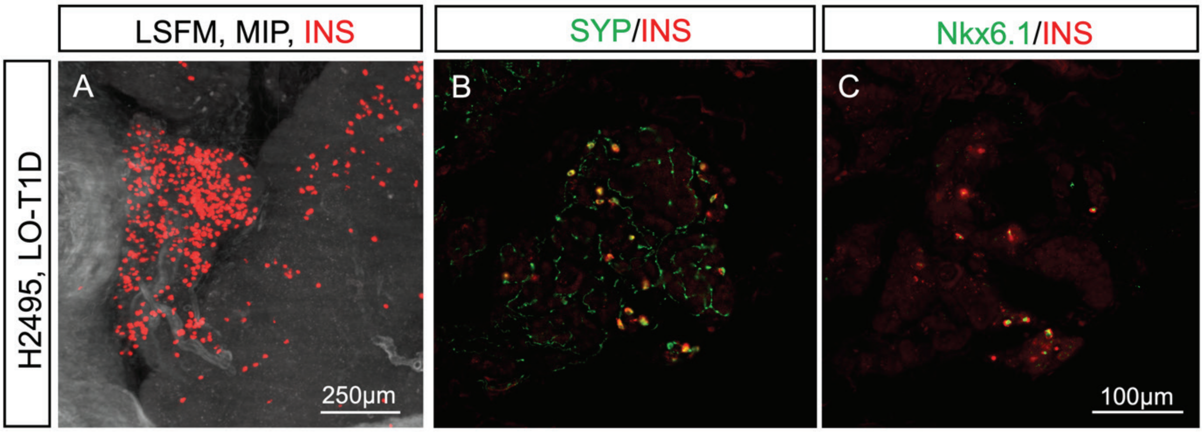


**Figure S6. Clustered extra-islet β-cells is a feature of LO-T1D pancreas. A,** LSFM MIP image showing a β-cell cluster from a LO-T1D pancreas (H2495, Male, 70 years, diagnosed at 4years, HbA1c 53.0). **B, C**, Slide scanner photomicrographs showing the β-cell cluster seen in (A), stained for INS (B and C, red) and Synaptophysin (B, green) and Nkx6.1 (C, green). Cluster of mature β-cells that are not intermingled by other endocrine cell-types could be detected in several additional pancreata.

| Metabolic state | ID | Sex | Age | BMI | HbA1c | Diabetes duration | Glucose Lowering therapy | COD | WIT | CIT |
|---|---|---|---|---|---|---|---|---|---|---|
| Type 1 Diabetes | H2442 | Male | 67 | 19.3 | 52 | 17 | Insulin | DBD | <5min | 5h 30min |
| Non-Diabetic | H2457 | Male | 28 | 23.7 | 35 | N.A. | N.A. | DBD | <5min | 2h |
| Non-Diabetic | H2522 | Female | 45 | 25.1 | 30 | N.A. | N.A. | DBD | <5min | 20h |

**Table S1. Clinical information of the pancreatic donors displayed in this study.** Abbreviations, COD: cause of death; WIT: warm ischemia time; CIT: cold ischemia time; DBD: donation after brain death.

### DESCRIPTION OF ADDITIONAL FILES

**Movie S1.** Maximum intensity projection (MIP) of combined NIR-OPT datasets displaying the complete β-cell mass distribution of a LO-T1D pancreas. The displayed pancreas (H2442) contains 0.02cm$^3$ INS$^+$ cells, comprising 1.73x 10$^5$ INS$^+$ objects at the current resolution. Note, due to size limitations the movie is significantly downsized.

**Movie S2**. Maximum intensity projection (MIP) from a LSFM scan of a representative ROI (see methods) from donor (H2442), stained for insulin (INS, red) and glucagon (GCG, green) showing examples of clustered and scattered extra-islet β-cells.
